# An atrial fibrillation-associated regulatory region modulates cardiac *Tbx5* levels and arrhythmia susceptibility

**DOI:** 10.1101/2022.05.14.491627

**Authors:** Fernanda M. Bosada, Karel van Duijvenboden, Mathilde R. Rivaud, Jae-Sun Uhm, Arie O. Verkerk, Bastiaan J. Boukens, Vincent M. Christoffels

## Abstract

Heart development and rhythm control are highly Tbx5 dosage-sensitive. *TBX5* haploinsufficiency causes congenital conduction disorders, whereas increased expression levels of *TBX5* in human heart samples has been associated with atrial fibrillation. We deleted the conserved mouse orthologues of two independent AF-associated genomic regions in the *Tbx5* locus, one intronic (*RE(int)^-/-^*) and one downstream of *Tbx5* (*RE(down)^-/-^*). In both lines we observed a modest (30%) increase of *Tbx5* in the postnatal atria. To gain insight into the effects of slight dosage increase *in vivo*, we investigated the atrial transcriptional, epigenetic and electrophysiological properties of both lines. We observed induction of genes involved in development, ion transport and conduction, increased action potential duration and increased susceptibility to atrial arrhythmias. We identified an AF-associated variant in the human intronic regulatory region that increases transcriptional activity. Expression of the AF-associated transcription factor *Prrx1* was induced in *RE(int)*^-/-^ cardiomyocytes. We found that some of the transcriptional and functional changes in the atria caused by increased *Tbx5* expression were normalized when reducing cardiac *Prrx1* expression in *RE(int)*^-/-^ mice, indicating an interaction between these two AF genes. We conclude that modest increases in expression of dose-dependent transcription factors, caused by common regulatory variants, significantly impact on the cardiac gene regulatory network and disease susceptibility.

## Introduction

The lifetime risk of developing a common disease, such as cardiovascular or neurodegenerative conditions, is influenced by genetic predisposition resulting from large numbers of inherited common genetic variants (single nucleotide polymorphisms; SNPs). Disease-associated variants are typically found in noncoding genomic regions and are thought to affect the functionality of regulatory elements (REs) such as enhancers or elements involved in chromatin conformation (1–5). These regulatory variants are pleiotropic and their effects on target gene expression are often specific to particular cell-types, conditions or stages of development (6–9). It remains challenging to identify the causal variants among the many associated variants in a disease-associated noncoding DNA region and the REs that are affected by such variants (10). Moreover, common variants typically have a small effect on phenotype, and the different functional variants may act additively, synergistically or oppositely. As a consequence, very few biological mechanisms linking disease-associated variant(s) to phenotype have been uncovered (11, 12) (13). Here, we set out to investigate how particular noncoding regions harboring clustered variants associated with a common disease modulate expression of a disease-associated transcription factor gene in a tissue-specific manner, and how this expression change affects phenotype *in vivo*.

Genome-wide association studies (GWAS) have identified many common variants in over 100 genetic loci associated with atrial fibrillation (AF) risk, the most prevalent arrhythmia associated with high comorbidity and increased mortality risk (14–19). The identification of functional variants, REs, and target genes underlying AF will provide important insights into the molecular mechanisms of disease (20). AF-associated variants have been identified in loci harboring transcription factor-encoding genes, including *PITX2, TBX5* and *PRRX1*, suggesting that altered expression levels of such factors cause imbalances in gene regulatory networks that control heart rhythm and function (20). Indeed, using mouse models, insufficiency of these transcriptional regulators was shown to cause arrhythmia susceptibility (21–29). Haploinsufficiency or heterozygous missense mutations in *TBX5* cause Holt-Oram syndrome in humans, characterized by forelimb malformations, congenital heart defects, and cardiac conduction anomalies, as a result of profound changes in gene regulatory networks in mouse and human cell models (30–35). Interestingly, a gain-of-function pathological missense variant in *TBX5* causes paroxysmal AF (36, 37). Moreover, a previous study uncovered a slight increase, rather than a reduction, in cardiac *TBX5* expression in human heart tissues has been associated with AF (19). The effects of small dosage increase in transcriptional regulators such as TBX5 are not well characterized.

We deleted the mouse orthologues of two AF variant-rich regions in the human *TBX5* locus to investigate how variant regions associated with a common disease modulate phenotype in a tissue- and developmental stage-specific manner. Each deletion caused a modest 30% increase in *Tbx5* expression in different heart compartments and at different stages of life. We report the relatively large effect of this modest increase in *Tbx5* expression on atrial function including arrhythmia susceptibility, and on the gene regulatory network. Decreased expression of *Prrx1* has been associated with AF in human and mouse models (24, 25). We observed a genetic interaction between *Tbx5* and *Prrx1*, and found that some of the transcriptional and functional changes in the atria caused by increased *Tbx5* expression were rescued by reducing cardiac *Prrx1* expression.

## Results

### Identification of two AF-associated regulatory regions in the *TBX5* locus

The topologically associated domain (38, 39) harboring *Tbx5* shows very limited contact with the adjacent domains harboring *Tbx3* and *Rbm19*, respectively (40). Because all AF-associated variants in this locus are found within the topologically associated domain of *TBX5* (Figure 1A), we anticipate that REs affected by the risk variants modulate the expression of *TBX5* only. Promoter capture Hi-C maps from iPSC-derived cardiomyocytes (41) show distinct contacts between the promoter of *Tbx5* and distal AF-associated regions (Figure 1A). To identify candidate REs we analyzed epigenomic datasets in both human (42, 43) and mouse orthologous region (Figure 1A-B). We selected two regions: the first situated in the last intron (RE(int)), and the second immediately downstream (RE(down)) of *Tbx5*, both containing highly conserved fragments which overlap with accessible chromatin sites in the left atria and ventricles (43, 44), cardiac H3K27ac and H3K4me1 ChIP-seq signatures (45), and EMERGE enhancer prediction signal (43, 44) (Figure 1B). The SNPs in the last intron and those downstream of the gene are clustered into two distinct haplotypes, suggesting that these two regions are independently inherited (Supplemental figure 1) (46). Using CRISPR/Cas9 genome editing, we deleted these candidate REs from the mouse genome to test their function *in vivo*.

**Figure 1.**
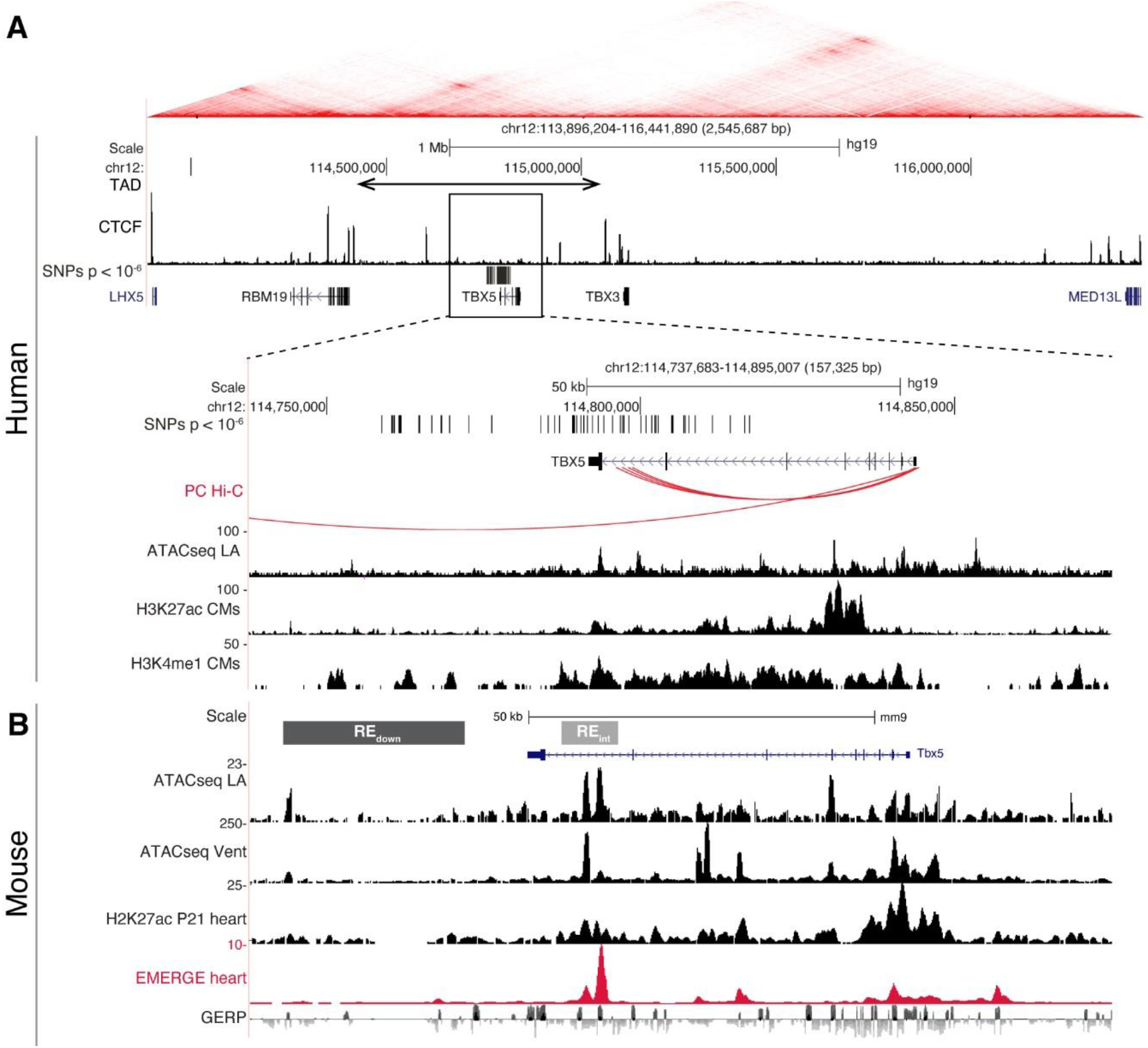
AF-associated noncoding variants are found in the *TBX5* locus. (A) Hi-C heatmap from human lymphoblastoid line GM12878 shows AF-associated variants are found in the regulatory domain of *TBX5*. Zoom-in of the AF-associated region overlaid with promoter capture Hi-C (red arcs), regions of open chromatin in whole left atria (ATACseq LA), H3K27ac and H3K4me1 ChIPseq signatures in cardiomyocytes. (B) Mouse orthologue of the human region including ATACseq from left atrial and ventricular CMs, H3K27ac ChIPseq from whole juvenile hearts, EMERGE, and conservation tracks. CRISPR/Cas9-generated deletions in light grey (REint), and dark grey (REdown).

### *Tbx5* expression and arrhythmia predisposition are increased in atria of *RE(int)*^-/-^ and *RE(down)*^-/-^ mice

Before birth, expression of *Tbx5* in the atria was not different between genotypes (Supplemental figure 2A). However, in juvenile atria of *RE(int)*^-/-^ mice (p=0.001), *Tbx5* expression was slightly increased compared to controls (Figure 2A). Expression of *Rbm19, Tbx3* and *Med131*, which neighbor *Tbx5*, remained unchanged in atria or ventricles of both mutants (Supplemental figure 2B). Both atria of *RE(int)*^-/-^ adult mice expressed approximately 30% more *Tbx5*, whereas only the left atrium of *RE(down)*^-/-^ adult mice expressed more *Tbx5* (Figure 2B-C). Interestingly, eQTL analysis shows that AF is associated with slightly increased *TBX5* expression in cardiac tissue (19). This suggests that AF-associated variants in the corresponding human regions may mediate the increase in *TBX5* expression observed in patients with risk variants.

**Figure 2.**
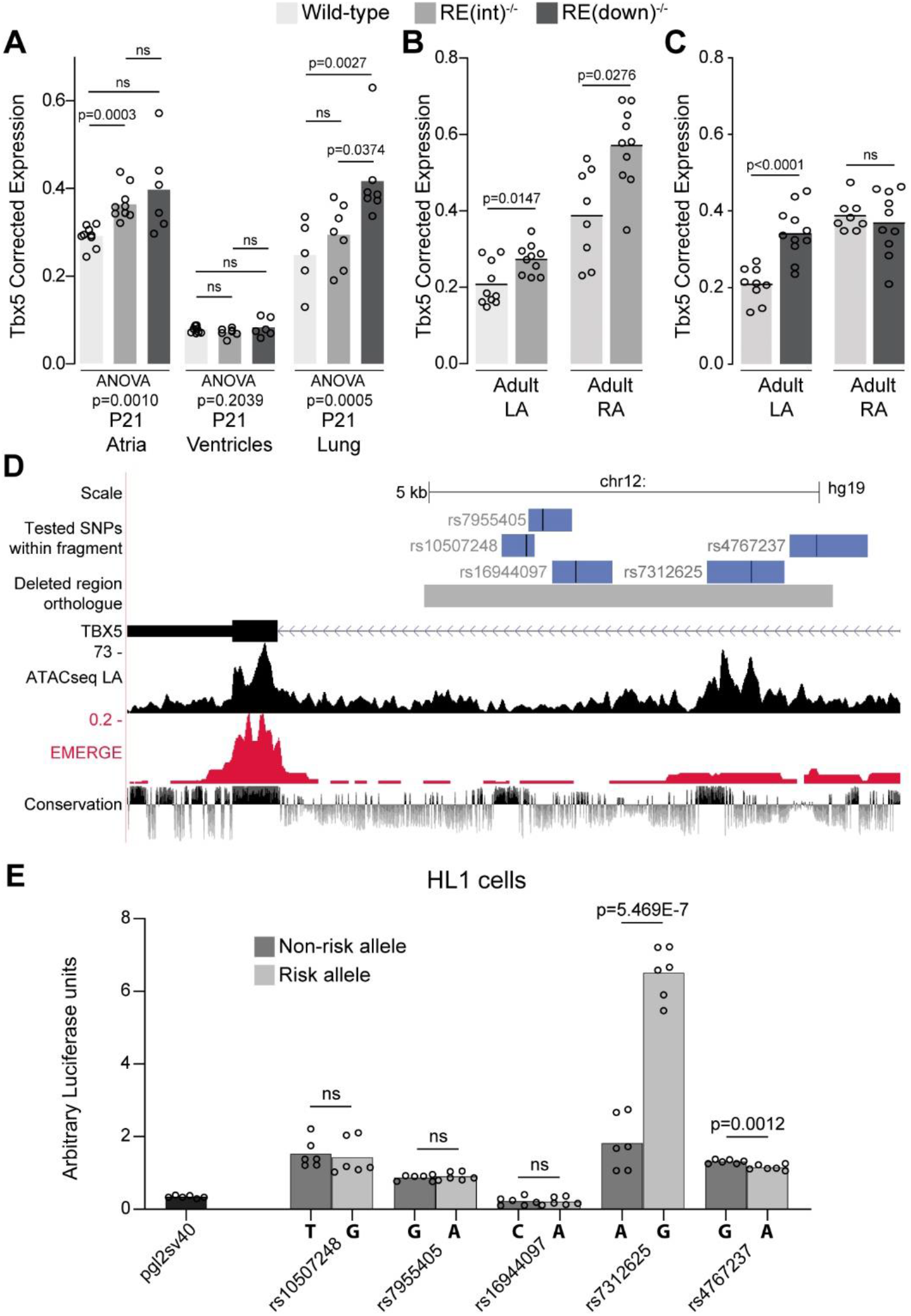
Deletion of AF-associated regions results in increased *Tbx5* in adult atrial tissue. (A) *Tbx5* expression in atria, ventricles, and lungs from P21 control, *RE(int)*^-/-^, and *RE(down)*^-/-^ mice determined by RT-qPCR. (B-C) *Tbx5* expression levels in adult control and *RE(int)*^-/-^ (B) or *RE(down)*^-/-^ (C) left and right atria. (D) UCSC browser view of the human intronic region (grey) and tested fragments containing AF-associated SNPs (blue) overlaid with chromatin conformation, EMERGE, and conservation tracks. (E) Luciferase assay (n=6) shows enhancer activity differences between non-risk (dark grey) or risk (grey) alleles (Kruskal-Wallis p=0.0019). Statistical significance within each tissue type was determined with ANOVA followed by pairwise comparisons using Dunnett’s T3 multiple comparison test in A, unpaired t-tests in B, C, and Kruskal-Wallis tests followed by pairwise unpaired t-tests (p values shown in figure) in E.

To identify functional AF-associated variants within the human intronic RE region, we tested enhancer activity of 5 fragments containing an AF-associated SNPs (p<10^−6^) (19) using luciferase assays in the atrial cardiomyocyte-like cell line HL-1 (Figure 2D). Of the tested fragments, the fragment containing rs7312625 showed increased enhancer activity, and the fragment containing rs4767237 showed minimal decreased activity (Figure 2E).

Next, we recorded *in* vivo electrocardiograms (ECGs) to determine the functional consequences of such a modest increase in *Tbx5*. Both *RE(int)*^-/-^ and *RE(down)*^-/-^ mice had slower and more variable heart rates (RR; SDNN) (Figure 3A, B). Additionally, we detected instances of sinus pauses and inverted P waves in mice with both deletions (Supplemental figure 3A). PR interval was significantly increased in *RE(int)*^-/-^ mice, but remained unaffected in *RE(down)*^-/-^ mice (Figure 3C). Other ECG parameters were not affected (Supplemental figure 3B). Heart rate corrected sinus node recovery time (cSNRT) and Wenckebach cycle length (WBCL) measured during transesophageal pacing (25, 47) were increased in both mouse models (Figure 3D, E), suggesting altered conduction system function. We next tested whether AA could be induced in both mouse lines using *in vivo* burst pacing (Figure 3G (typical AA traces)). The total duration of all AA episodes per mouse was greater in both deletion models when compared to control littermates (Figure 3F (top graph)). Yet, we were able to induce AAs more often in *RE(int)*^-/-^, but not *RE(down)*^-/-^ mice compared to controls (Figure 3F (below graph)).

**Figure 3.**
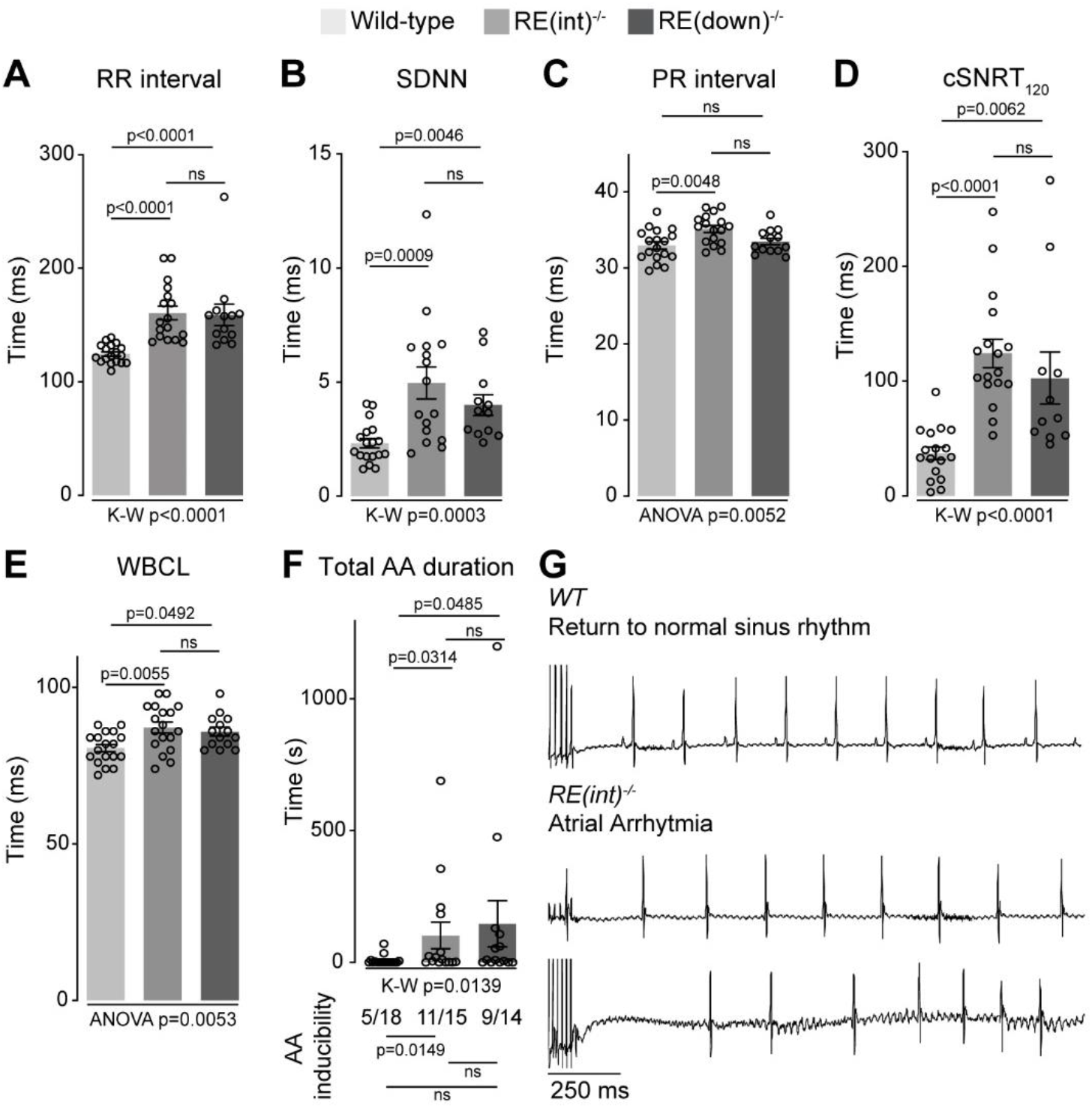
Increased *Tbx5* in adult atria results in altered *in vivo* electrophysiology. (A-C) Graphs show individual and average ECG measurements for heart rate (RR) (A), heart rate variation (SDNN) (B), and PR-interval (C) of adult *wild-type, RE(int)*^-/-^, and *RE(down)*^-/-^ mice. (D, E) Graphs show changes in heart rate-corrected sinus node recovery times at 120 ms (cSNRT_120_) (D), and Wenckebach cycle length (WBCL) (E). (F) Bar graph depicts the total time each mouse spent in an atrial arrhythmia (AA) episode after two pacing passes, with the number of mice in which at least one episode lasting >1sec was observed below each bar. (G) Representative traces from *wild-type* (top) and two *RE(int)*^-/-^ (bottom) individuals showing disappearance of p waves or the start of atrial arrythmia with variability in ventricular response after pacing stimulus. Significance for *in vivo* parameters was determined with Kruskal-Wallis test followed by Dunn’s multiple comparison tests in A, B, D, and AA duration in F (top graph), and one-way ANOVA followed by Tukey’s multiple comparisons test in C and E. AA inducibility significance was determined with pairwise Fisher’s exact test (F bottom of graph).

To further characterize the electrophysiological phenotypes of *RE(int)*^-/-^ mice we analyzed the properties of single isolated left atrial cardiomyocytes using patch-clamp. At 6 Hz stimulation, we observed APD prolongation in mutants at all measured repolarization stages, with no changes in AP upstroke velocity, resting membrane potential and maximal AP amplitude (Figure 4A-C). APD increase was present at all frequencies measured (Figure 4D). Because *Tbx5* has important roles in intracellular Ca^2+^ handling (26, 27), we measured intracellular Ca^2+^ concentration ([Ca^2+^]_i_) in isolated atrial CMs using fluorescent calcium indicator Indo-1. We did not observe any changes in systolic or diastolic [Ca^2+^]_i_ concentration (Figure 4E). Together, our data show that a slight increase in *Tbx5* disturbs atrial function and can predispose to arrhythmia.

**Figure 4.**
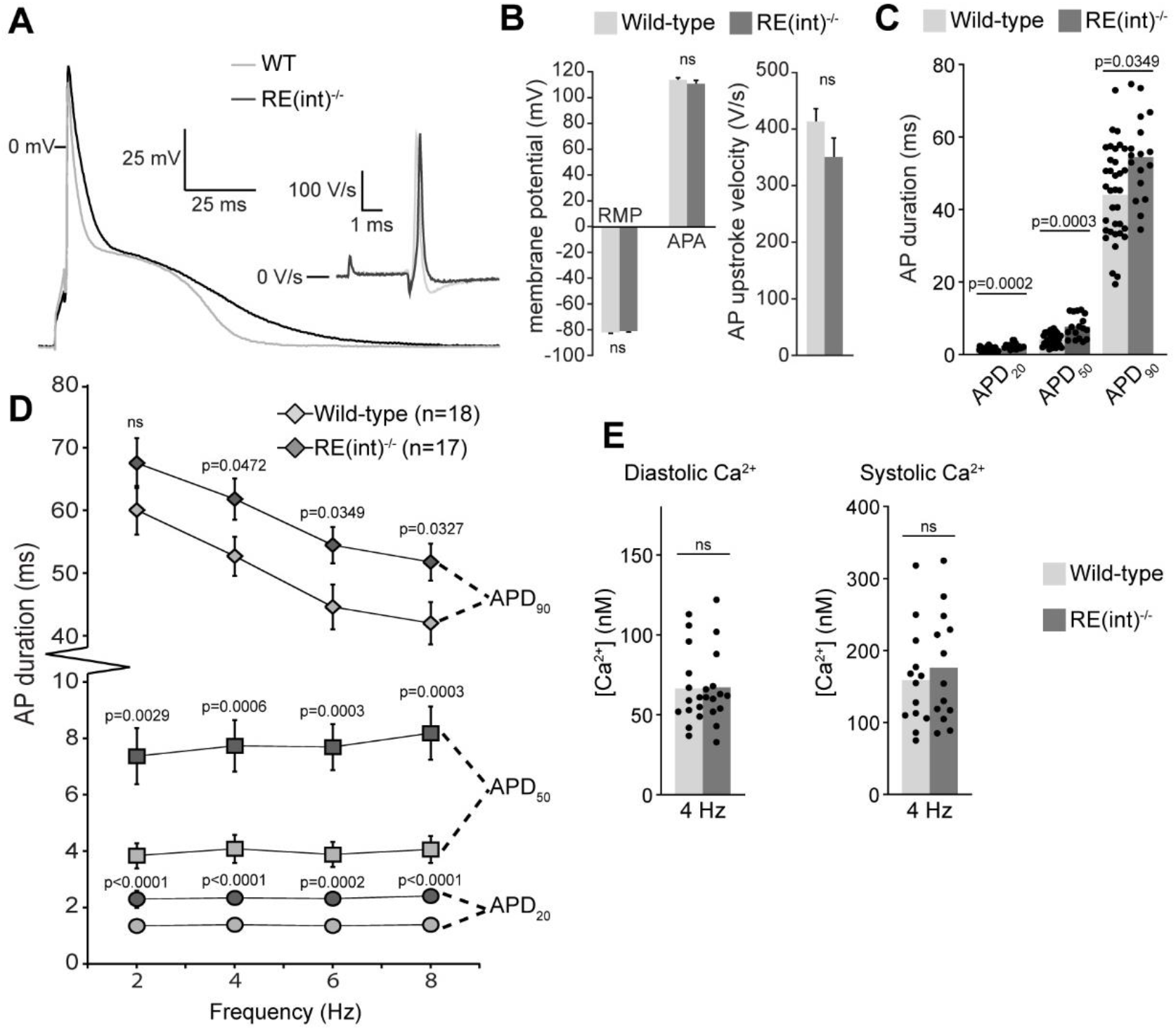
Atrial function abnormalities in *RE(int)*^-/-^ mice. (A) Typical action potentials (APs) and upstroke velocity measured at 6 Hz in single left atrial cardiomyocytes using the amphotericin-perforated patch clamp technique. (B) Average resting membrane potential (RMP), AP amplitude (APA), and AP upstroke velocity remain unchanged in mutants. (C) AP duration at 20, 50, and 90% of repolarization (APD_20_, APD_50_, APD_90_, respectively) is increased in mutants. (D) APD_90_ was prolonged at all tested frequencies in mutant left atrial cardiomyocytes. Error bars are SD. (E) Diastolic and systolic intracellular Ca^2+^ concentrations ([Ca^2+^]_i_) were not changed in *RE(int)*^-/-^ compared to controls. Statistical significance in B was determined with unpaired t-tests with Welch’s correction, and unpaired t-tests were used in B. Experimental groups were compared using two-way repeated measures ANOVA (D, E), followed by pairwise unpaired t-tests (C). Error bars are SEM.

### Sensitivity of the atrial gene regulatory network to modestly increased *Tbx5* dosage

To gain insight into the mechanism underlying the changes in electrophysiological properties in mutants, we performed transcriptional profiling of whole left atria. We detected 13790 and 13564 different transcripts in *RE(int)*^-/-^ and *RE(down)*^-/-^, respectively, of which 816 were significantly up- and 1235 were downregulated in *RE(int)*^-/-^ (Figure 5A; adjusted p<0.05), and 112 were up- and 123 were downregulated in *RE(down)*^-/-^ (Figure 5B; adjusted p<0.05). Both datasets shared a high proportion of deregulated genes, with fewer significantly deregulated genes observed in *RE(down)*^-/-^ compared to *RE(int)*^-/-^ (Figure 5C). These data indicate that transcriptomes in *RE(int)*^-/-^ atria and in *RE(down)*^-/-^ atria are comparably affected qualitatively, but not quantitatively, consistent with the slightly higher increase in *Tbx5* expression in *RE(int)*^-/-^ compared to *RE(down)*^-/-^. Gene Ontology analysis (48) revealed that processes involved in cellular compartment organization, ion transport, and cardiac conduction characterized the transcripts found in the upregulated genes, and extracellular matrix organization, vasculature development, and actin cytoskeleton organization were found in the downregulated set in both deletions (Figure 5D). Accordingly, several *Tbx5* target genes known to affect APD or whose deregulation may result in electrophysiological changes, (21, 26, 27) were significantly deregulated in one or both deletion lines (Figure 5F, Supplemental Figure 4A).

**Figure 5.**
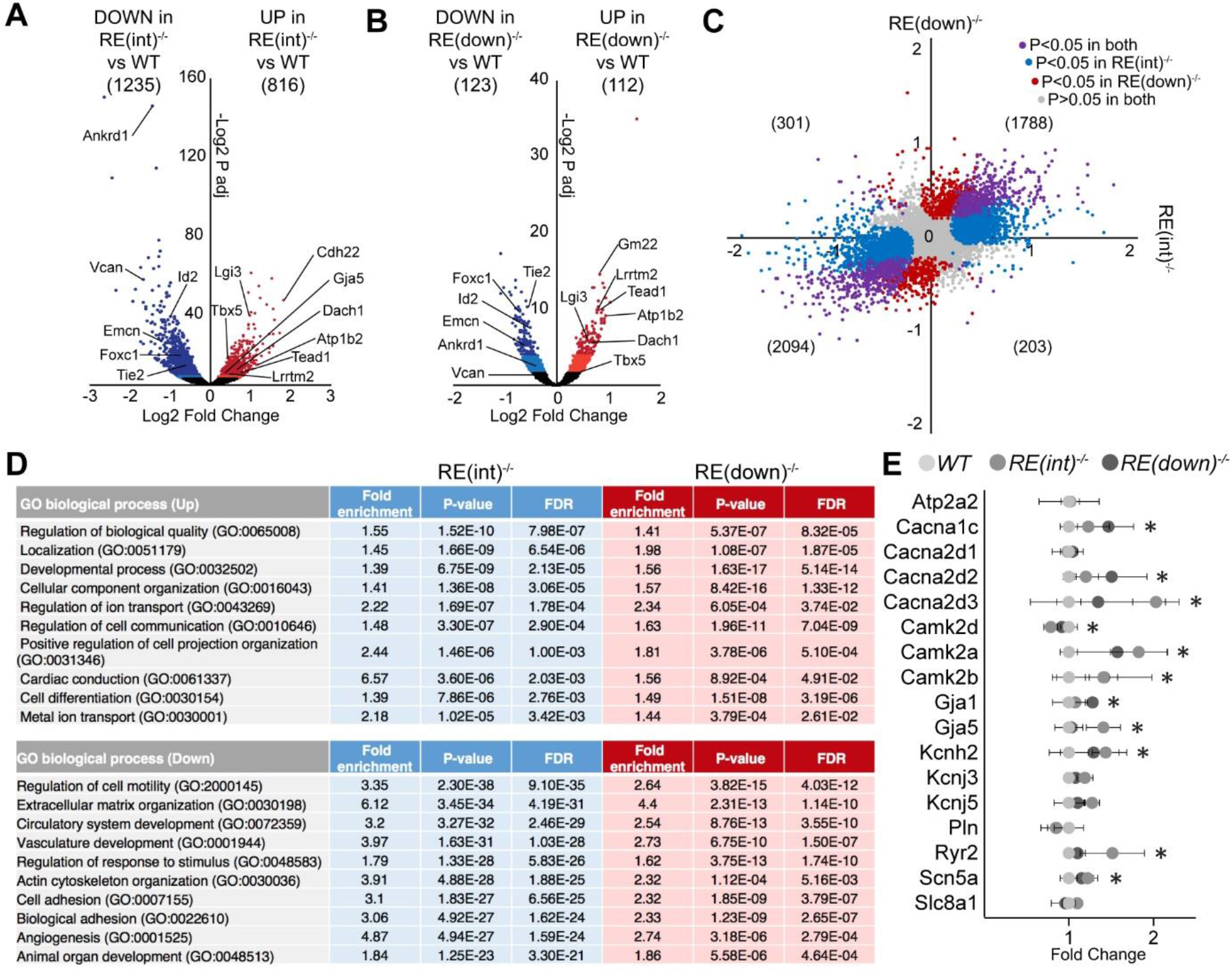
Transcriptomic analysis of *RE(int)*^-/-^ left atria. (A, B) Volcano plot showing differentially expressed transcripts in *RE(int)*^-/-^ (n=3)(A) and *RE(down)*^-/-^ (n=3) (B) from *wild-type* (n=4) left atria. Dark red and light red dots indicate significantly upregulated genes by raw p-value and p-adjusted for multiple testing, respectively. Correspondingly, dark blue and light blue dots indicate downregulated genes. P values were adjusted for multiple testing using the false discovery rate (FDR) method of Benjamini-Hochberg. (C) X-Y plot of all transcripts in *RE(int)*^-/-^ (x axis) and *RE(down)*^-/-^ (y axis), with deregulated genes common to both deletion mutants in purple, *RE(int)*^-/-^ deregulated genes in blue, and *RE(down)*^-/-^ deregulated genes in red. (D) Gene ontology (GO) analysis of upregulated and downregulated genes in *RE(int)*^-/-^ and *RE(down)*^-/-^ mutants. (F) Graph depicts fold change expression in control and mutant samples of genes known to affect action potential duration. * denotes significantly deregulated in one or both mutant lines.

To determine whether a slight increase in *Tbx5* expression in cardiomyocytes results in changes in chromatin accessibility, we performed ATAC-sequencing profiling (ATACseq) of *RE(int)*^-/-^ left atrial cardiomyocytes (49). After peak-calling, we found a total of 85,569 accessible sites common to both genotypes, and only 15 sites of increased accessibility and 32 of decreased accessibility in mutants (Supplemental Figure 5A). Two of the sites with decreased accessibility were found in the locus of *Tbx5* (one of which caused by the deletion of the intronic genomic fragment in *RE(int)*^-/-^ mice), suggestive for increased direct transcriptional autoregulation in *RE(int)*^-/-^ mice (Supplemental Figure 5B). The very minor changes in chromatin accessibility indicate that a modest increase in *Tbx5* expression does not significantly change transcription factor occupancy or epigenetic state.

### An interaction between *Tbx5* and *Prrx1*, two AF-associated genes

We noted that *RE(int)*^-/-^ cardiomyocytes express more *Prrx1*, encoding the transcription factor Paired Related Homeobox 1. Common variants in the *PRRX1* locus have been associated with AF, and a cardiac-specific decrease in *PRRX1* expression has been linked to AF-predisposition (19, 24, 25). *Prrx1(enh)*^-/-^ mice express less *Prrx1* specifically in cardiomyocytes compared to controls, and show atrial conduction slowing, lower AP upstroke velocity (indicative for lower Na^+^ current densities)(50), as well as increased systolic and diastolic [Ca^2+^]_i_ concentration that culminate in increased susceptibility to atrial arrhythmia induction (25). To explore whether and how these two AF-risk genes may interact, we intercrossed *RE(int)*^-/-^ (increased atrial *Tbx5*, increased *Prrx1* in cardiomyocytes) with *Prrx1(enh)*^-/-^ mice (decreased *Prrx1* in cardiomyocytes), and investigated cardiac transcriptomes and phenotypes across genotypes. We isolated cardiomyocyte and non-cardiomyocyte nuclei from whole hearts using a PCM-1 antibody and interrogated *Tbx5* and *Prrx1* expression across all genotypes. *Tbx5* was upregulated in *RE(int)*^-/-^ cardiomyocytes, as expected, but not deregulated in *Prrx1(enh)*^-/-^ or double mutant cardiomyocytes (Figure 6A). *Prrx1* levels were increased in *RE(int)*^-/-^ cardiomyocytes and decreased in *Prrx1(enh)*^-/-^, as expected, and also decreased in double mutant cardiomyocytes (Figure 6B). There were no statistically significant changes detected in the non-cardiomyocyte fractions (Figure 6A, B). These data indicate that in cardiomyocytes, Tbx5 regulates *Prrx1*, and that a regulatory feedback loop modulates *Tbx5* levels when *Prrx1* expression is reduced due to the *Prrx1(enh)* deletion.

**Figure 6.**
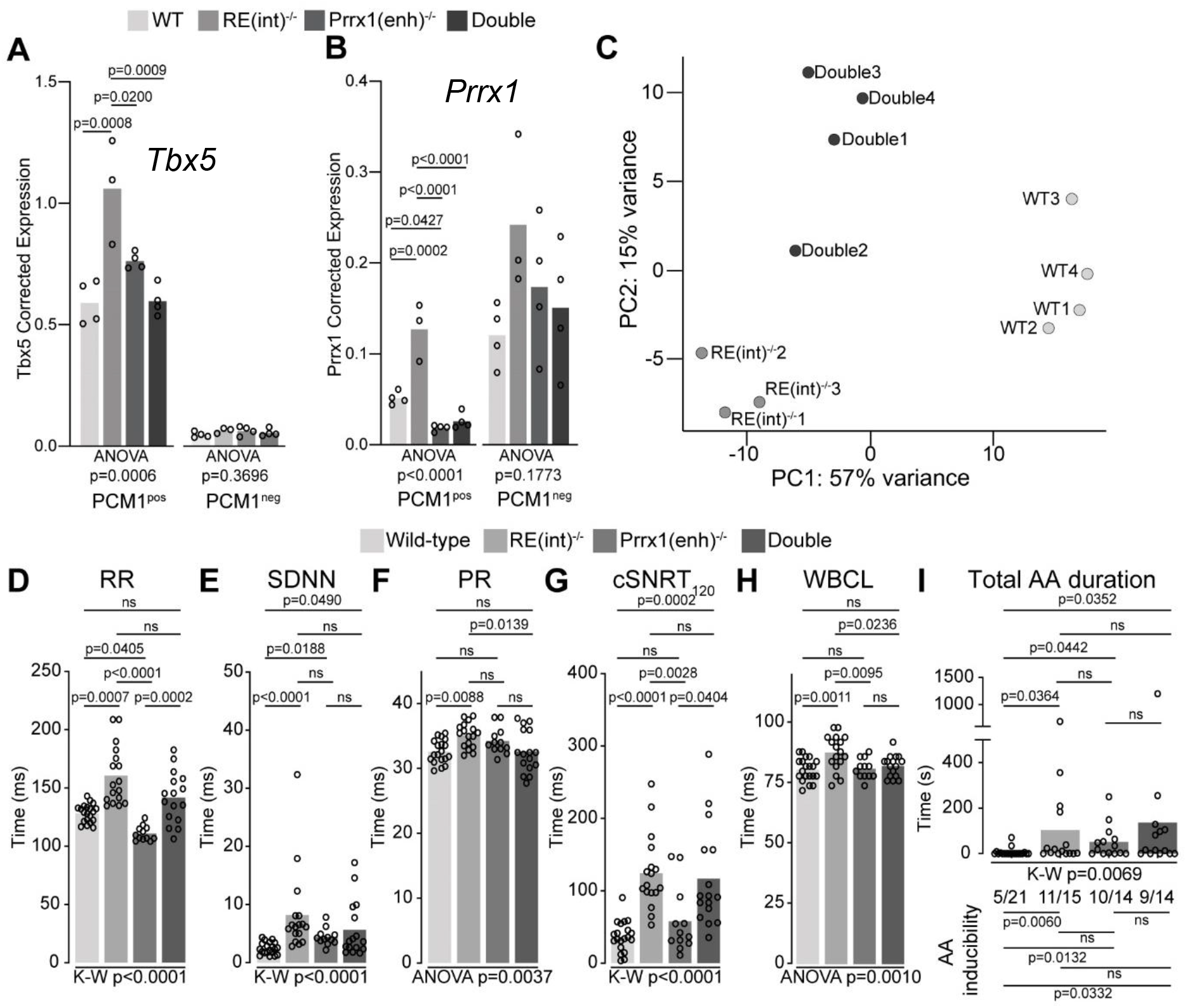
A genetic interaction between *Tbx5* and *Prrx1*. (A, B) Graph shows reference gene-corrected expression of *Tbx5* (A), and *Prrx1* (B) in PCM1-positive and -negative nuclei fractions of *wild-type, RE(int)^-/-^, Prrx1(enh)*^-/-^ and *double* mutant whole hearts determined by RT-qPCR. (C) Principal component analysis of transcriptomes of *wild-type, RE(int)*^-/-^ and *Double*^-/-^ left atrial samples. (D-H) Graphs show individual and average ECG measurements for RR (D), heart rate variation (SDNN) (E), PR interval (F), cSNRT at 120 ms pacing (G), and WBCL (H) of *wild-type, RE(int)^-/-^, Prrx1(enh)*^-/-^ and *Double*^-/-^ mice. (I) Bar graph depicts the total time each mouse spent in an AA episode after two pacing passes, with the number of mice in which at least one episode lasting >1sec was observed below each bar. Statistical significance was determined using one-way ANOVA followed by Tukey’s test for pairwise comparisons in A, B, F, H, Kruskal-Wallis test followed by Dunn’s test for pairwise comparisons in D, E, G, and AA duration in I (top of graph), Fisher’s exact test for pairwise comparisons of AA inducibility in I (bottom of graph).

We next performed transcriptome analysis and found that deregulated genes in left atria of *RE(int)^-/-^; Prrx1(enh)*^-/-^ double mutants were more similar to the transcript changes found in *RE(int)*^-/-^ than the ones found in *Prrx1(enh)*^-/-^, when comparing all lines to WT, suggesting a dominant contribution of *RE(int)*^-/-^ to the transcriptional changes in double mutants (Supplemental Figure 6A, B). Using principal component analysis, we found that transcriptomes clustered according to genotype (Figure 6C). Compared to atrial transcriptomes of *RE(int)*^-/-^ mice, those of *RE(int)^-/-^; Prrx1(enh)*^-/-^ double homozygous showed less variance with WT atrial transcriptomes (Figure 6C, Supplemental Figure 6). This suggested an interaction between *Tbx5* and *Prrx1*, in which reduced *Prrx1* expression partially normalizes increased Tbx5-induced transcriptomic changes.

Next, we considered whether the adult electrophysiological changes observed in *RE(int)*^-/-^ would also be rescued by the presence of this second AF-risk allele. *In vivo* ECGs and burst pacing experiments revealed that RR, PR interval, and WBCL were normalized in double homozygous mice, whereas SDNN, cSNRT, and AA inducibility remained unchanged (Figure 6D-I).

Taken together, our findings suggest that an interaction between *Tbx5* and *Prrx1* exists; in which *Prrx1* expression is induced by Tbx5, where *Prrx1* is required to induce *Tbx5* expression in *Prrx1(enh)*^-/-^ mice, and increased Tbx5 imposes changes in expression and electrophysiological properties, in part through increasing *Prrx1* expression in cardiomyocytes.

## Discussion

Our study reveals that mouse orthologues of two independent variant regions in the *TBX5* locus modulate *Tbx5* expression in tissue- and stage-specific manners, causing distinct specific phenotypes. While the impact of 50-100% dose reduction of *Tbx5* in humans and model systems has been well-investigated (33–35, 51–54), we here show the effect of physiologically relevant increases in expression of *Tbx5*. This is significant as both decreased and increased expression of genes has been associated with increased AF risk.(19). Our study reveals that atrial- and postnatal-specific increase in *Tbx5* levels of only 30% affects postnatal atrial gene regulation, function and arrhythmia propensity. Furthermore, we provide an example of the interaction between the effect of two independent AF-associated variant regions (*TBX5* and *PRRX1)* on phenotype. Our models provide insight into the mechanisms underlying the pleiotropic effects and interactions of disease-associated regulatory variants, which typically cause small changes in target gene expression in particular cell-types, conditions or stages of development (6–9).

We found that the mouse orthologues of two noncoding regions studied here containing AF-associated variants in the *TBX5* locus harbor REs that independently controlled *Tbx5* expression in the heart *in vivo*. Deletion of either of the two RE-containing regions only affected *Tbx5* expression, consistent with the notion that *Tbx5* and surrounding RE regions are largely confined to one topologically associated domain not shared with adjacent genes (40). The mechanism of the repressive action of the REs and cross-talk with the other REs in the locus (55) remain to be established. For *RE(int)*, which showed the largest effect size in atria upon deletion, we identified two AF-associated SNPs that caused differential transcriptional activity of the RE sub fragment in HL1 atrial cardiomyocyte-like cells; one variant caused ~6 fold increase and the other minimally decreased activity. We speculate that in individuals carrying the first, activity-increasing variant, the repressive activity of *RE(int)* is reduced in the context of the entire regulatory system, thus causing increased *TBX5* expression. The opposite could be true for the second variant, yet the small effect would likely be experimentally undetectable.

Heart rates of the *RE(int)*^-/-^ mice were slower and more variable than of controls. In addition, PR interval was prolonged in *RE(int)*^-/-^ mice combination with increased Wenckebach cycle lengths. The latter is in line with the important role of Tbx5 in sinus node and AV node development and function (33, 34, 56, 57). Moreover, atrial arrhythmias could be more easily induced in *RE(int)*^-/-^ mice suggesting electrophysiological remodeling of the atria. It has been shown that mice haploinsufficient for *Tbx5* show atrial downregulation of genes encoding proteins involved in cardiac conduction (e.g. *Gja1/Cx43* and *Scn5a*/Nav1.5) and Ca^2+^ handling (e.g *Atp2a2*/Serca2a and *Ryr2*/Ryr2) (21). This leads to slower conduction, reduced Ca^2+^ concentration in the sarcoplasmic reticulum [Ca^2+^]SR, increased incidence of early and delayed afterdepolarization, and accordingly, increased AF propensity (21, 26). In the atria of our *RE(int)*^-/-^ mice, where expression of *Tbx5* was slightly elevated, transcriptomic analysis indicated upregulation of genes involved in cardiac conduction including *Scn5a* and *Cx43*. The Ca^2+^ handling genes *Atp2a2*/Serca2a and *Ryr2*/Ryr2 were not differently expressed between *RE(int)*^-/-^ mice and controls. Accordingly, in our mice, we did not observe prolonged Ca^2+^ transients and long decay constants as seen in *Tbx5* haploinsufficient mice (26). Expression of *Cacna1c* and *Cacna2d2/Cacna2d3*, encoding subunits of the L-type Ca^2+^ channel, was upregulated in the atrium of *RE(int)*^-/-^ mice. However, diastolic calcium concentration was not different between mice indicating that the L-type Ca^2+^ current (I_Ca,L_) was not affected in *RE(int)*^-/-^ mice. We did observe longer atrial APD in *RE(int)*^-/-^ mice compared to controls, which may predispose to increased arrhythmia inducibility, as seen in mice harboring the pathogenic TBX5-G125R variant or *Prrx1* enhancer deletion (25, 37). Although shortened APD is typically found in AF patients with sustained arrhythmia (58, 59), prolonged APD has been previously associated with increased risk of developing AF (25, 37, 60–62).

Not all aspects of the molecular mechanism underlying electrophysiological remodeling have become clear so far. We speculate that the observed modest changes in gene expression led to proportional altered electrophysiological properties that are below experimentally detectable thresholds. Furthermore, previous research has demonstrated that Tbx5 and Pitx2 work antagonistically in the left atrium to tightly regulate the expression of genes impacting on cardiac electrophysiology (21). Yet, we did not detect a change in *Pitx2* expression in the atria of *RE(int)*^-/-^ mice. Nevertheless, our data showed that perturbation of this regulatory network in either up or down direction can lead to increased arrhythmia susceptibility.

The risk of developing complex diseases such as AF is strongly influenced by a large number of pleiotropic variants that each confers a small change in overall risk. We found that Tbx5 drives *Prrx1* expression, and that *Tbx5* levels are Prrx1-dependent (or Prrx1 target-dependent) in specific contexts. We then asked whether combining two alleles modeling the changes in gene expression conferred by AF-associated variant regions would cause a greater effect on heart phenotype, and by extension, arrhythmia predisposition, than each allele alone. Reduced *PRRX1* expression has been associated with AF (19). We recently characterized a mouse model with a deletion of the orthologue of the region associated with AF in human causing reduced *Prrx1* expression, perturbation of atrial conduction and increased AF susceptibility (25). Here, we found that introduction of this AF-susceptibility allele rescued heart rate, PR interval, and atrioventricular node phenotypes detected in *RE(int)*^-/-^ mice. This example shows that two variant regions that are both independently associated with increased risk for a particular disease, do not necessarily cumulatively increase disease predisposing phenotype, but may neutralize each other’s effect. Previously, the atrial electrophysiological phenotype of Tbx5 haploinsufficient mice was observed to be rescued by haploinsufficiency of either *Pitx2* or *Gata4*, both of which have been associated with AF as well (21, 26). While the direction of change in these models does not necessarily correspond to the direction of change in AF patients, this highlights the inherent robustness of the underlying gene regulatory networks, which in general remain stable when individual quantitative parameters such as transcription factor dose or binding sites affinity change(63). This also implies that evaluating specific interactions between AF risk loci will be necessary for ascertaining individual risk from genetic association data.

In conclusion, our data provide unique mechanistic insights into the biological effects of variant region-driven modest physiologically relevant changes in expression of crucial transcriptional regulators, and into the impact of interactions between transcriptional output-modulating risk loci on atrial biology.

## Materials and Methods

A detailed Methods section is available in the Data Supplement.

## Ethics statement

Housing, husbandry, all animal care and experimental protocols were in accordance with guidelines from the Directive 2010/63/EU of the European Parliament and Dutch government. Protocols were approved by the Animal Experimental Committee of the Amsterdam University Medical Centers. Animal group sizes were determined based on previous experience.

## Data availability

Adult left atrial RNAseq and ATACseq have been deposited under GEO accession numbers GSE189342 and GSE189498.

## Statistics

The experimenters were blind to mouse genotype during all measurements and outcome assessment. Datasets were tested for normality using Shapiro-Wilk test unless specified otherwise. Whole tissue RT-qPCR fold change vs WT in fetal ventricles and adult whole left atria corrected expression changes were analyzed with unpaired t-tests and reference gene-corrected *Tbx5* in juvenile whole tissue was analyzed with Welch’s ANOVA followed by Dunnet’s T3 multiple comparison tests within each tissue type. Luciferase assays on HL1 cells were analyzed using Kruskal-Wallis test. *In vivo* electrophysiology was analyzed with Kruskal-Wallis followed by Dunn’s multiple comparison tests or one-way ANOVA followed by Tukey’s multiple comparison tests. Significance of atrial arrhythmia (AA) duration was determined with Mann Whitney Wilcoxon test and differences in AA inducibility were tested using Fisher’s exact test. Conduction velocity was analyzed with unpaired t-tests with Welch’s correction. For single cell and whole tissue action potential (AP) duration (APD) normality and equal variance assumptions were tested with the Kolmogorov-Smirnov and the Levene median test, respectively. Two groups were compared with unpaired t-test or repeated measures ANOVA followed by pairwise comparison using the Student-Newman-Keuls test. Differences in Ca^2+^ transient amplitude as well as changes in diastolic and systolic Ca^2+^ concentrations in atria were tested using two-way ANOVA. Multiple testing corrections were performed independently within each hypothesis. Data are presented as individual data points and mean or mean ± standard error of the mean (SEM) or standard deviation (SD), as indicated, and p<0.05 defines statistical significance. Statistical analysis was performed using GraphPad Prism 9.

## Acknowledgements

We thank Berend de Jonge for technical assistance. This work was supported by CardioVasculair Onderzoek Nederland (CVON) project 2014-18 CONCOR-genes Young Talent Program (to F.M.B.), CVON project 2014-18 CONCOR-genes (to V.M.C.), Leducq Foundation 14CVD01 (to V.M.C.), Dutch CardioVascular Alliance OUTREACH (to V.M.C.) and ZonMW TOP 91217061 (to V.M.C.).

AA: (atrial arrhythmia)
AF: (atrial fibrillation)
AP: (action potential)
cSNRT: (corrected sinus node recovery time)
RE: (regulatory element)
WBCL: (Wenckebach cycle length)

