## Supplemental Material for "An atrial fibrillation-associated regulatory region modulates cardiac *Tbx5* levels and arrhythmia susceptibility"

### Expanded materials and methods

#### Usage of epigenomic datasets

The following publicly available epigenomic datasets were used: Hi-C [1], AF variants [2], promoter capture Hi-C maps from iPSC-derived cardiomyocytes [3], accessible chromatin in human and mouse left atria [4], cardiac H3K27ac and H3K4me1 ChIP-seq signatures in human [5], and EMERGE enhancer prediction signal [4, 6].

#### Generation of mutant mice

Mutant mice were generated using CRISPR/Cas9. Guide RNA (sgRNA) constructs were designed with ZiFiT Targeter [7]. The sgRNA constructs were transcribed *in vitro* using MEGAshortscript T7 (Invitrogen AM1354) and mMessage Machine T7 transcription kit (Invitrogen AM1344) according to manufacturer instructions. One-cell FVB/NRj zygotes were microinjected with 10 ng/µL each sgRNA and 25 ng/µL Cas9 mRNA to generate mouse founders. Deletions were validated by PCR and Sanger sequencing. The sgRNA target sequences are the following: RE(int) guide 1 (GGGAAATCGCCTTACCTTTC), guide 2 (GGACTGTTGGGTCACCTTGT), and RE(down) guide 1 (GGCTCCTTCGTCAGTAAATA), guide 2 (AAACAAGGGCTCTCTGGCGTTT). Founders were backcrossed with wildtype FVB/NJ mice to obtain stable lines. Downstream experiments were performed on F3-F7 mice, backcrossed with wild-type FVB mice. All transgenic mice were maintained on a FVB/NJ background commercially obtained from Jackson laboratory (stock number 100800).

For tissue harvest, animals were euthanized by 20% $CO_2$ inhalation followed by cervical dislocation.

#### EdU cardiomyocyte proliferation assay

Timed pregnant female mice received intraperitoneal injection of 100 mg/kg EdU one hour before sacrifice by isoflurane and cervical dislocation. E14.5 fetuses were isolated in 1xPBS on ice, the head was removed and the body fixed in 4% PFA for 24 hours. 7 µm sections were stained with mouse-anti-actin (1:400; Sigma), goat-anti-nkx2.5 (1:150; Santa Cruz), DAPI (1:1000) and EdU Click-IT (ThermoFisher) before imaging and counting. Nkx2.5 positive nuclei were designated cardiomyocytes, Nkx2.5 and EdU double positive nuclei were designated proliferating cardiomyocytes. Ratios were normalized within each litter.

#### qPCR

Total RNA was isolated from atria, ventricle and lungs from P21 and left and right atria from adult male and female mice using ReliaPrep RNA Tissue Miniprep System (Promega, Z6112) according to the manufacturer’s protocol. cDNA was reverse transcribed with oligo dT primers from 500ng of total RNA, or random hexamers from 500pg of CM nuclear RNA, according to the manufacturer’s protocol of the Superscript II Reverse Transcriptase system

(Thermo Fisher Scientific, 18064014). Expression levels of candidate target genes were determined by quantitative real-time PCR using a LightCycler 480 Instrument II (Roche Life Science, 05015243001). Expression levels were measured using LightCycler 480 SYBR Green I Master (Roche, 04887352001) and the primers had a concentration of 1 pmol/L. The amplification protocol consisted of 5 minutes 95 °C followed by 45 cycles of 10 seconds 95°C, 20 seconds 60°C and 20 seconds 72°C. Relative start concentration (N0) was calculated using LinRegPCR [8]. Values were normalized to the geometric mean of two reference genes per experiment (Hprt, Ppia, or Rpl32) [9]. The primer sequences are as follows: (all 3' to 5') Hprt: TGTTGGATATGCCCTTGACT, GATTCAACTTGCGCTCATCT; Ppia: GGGTGGTGACTTTACACGCC, CTTGCCATCCAGCCATTCAG; Rpl32: GCCTCTGGTGAAGCCCAAG, TTGTTGCTCCCATAACCGATGT; Tbx5: CCCGGAGACAGCTTTTATCG, TGGTTGGAGGTGACTTTGTG; Prrx1: CACAAGCAGACGAAAGTGTGG, GTTGTCCTGTTTCTCCGCTG; Tbx3: CGCCGTTACTGCCTATCAGAA; GCCATTGCCAGTGTCTCGAA.

**Cell Culture and Transfection Luciferase Assays**

RE sub fragments were cloned into a modified pGL2-Basic plasmid containing an SV40 minimal promoter and an adjusted multiple cloning site for *in vitro* analysis by a transfection luciferase assay. HL1 cells were grown in 24-well plates in Claycomb medium (Sigma-Aldrich, 51800C) supplemented with chemically defined HL-1 FBS substitute (Lonza, 77227), Glutamax (ThermoFisher Scientific, 35050-061) and Pen/Strep (ThermoFisher Scientific, 15070-063). Cells were transfected using polyethylenimine 25 kDa (PEI, Brunschwig, 23966-2) at a 1:3 ratio (DNA:PEI). Standard transfections were carried out using 200 ng of reporter construct per well. 24 hours after transfection, cells were lysed using Renilla luciferase assay lysis buffer (Promega, E291A-C) and luciferase activity measured. Luciferase measurements were performed using a GloMax Explorer (Promega, GM3500). During the measurement, 100 µL D-Luciferin (p.j.k, 102111) was injected (150 µL/second) followed by a 1 second delay and 5 seconds of measurement. Transfections were carried out at least three times and measured in duplicates.

***In vivo* electrophysiology**

12-20 week old male mice were anesthetized with 5% Isoflurane (Pharmachemie B.V. 061756) and placed on thermostated mat (36ºC) with a steady flow of 1.5% isoflurane during all experiments. Electrodes were inserted subcutaneously in the limbs and connected to an ECG amplifier (Powerlab 26T, AD Instruments). The electrocardiogram (ECG) was measured for 5 minutes. ECG parameters were determined in Lead II (L-R) based on the last 60 seconds of the recording. For atrial stimulation, an octapolar CIB'er electrode (NuMED) was advanced through the esophagus to achieve atrial capture. Atrial capture thresholds were determined for each mouse, and all pacing protocols were performed at 2x threshold. For sinus node recovery time (SNRT) measurements, a 4 second pacing train with a cycle length of 120 or 100 ms was used. SNRT was defined as the interval between the last pacing stimulus and onset of the first P wave. To control for differences in sinus rate,

SNRT was normalized to resting heart rate (cSNRT = SNRT – RR interval). To determine the Wenckebach cycle length (WBCL) we applied a 4-second pacing train starting at a cycle length of 100 ms, and decreasing by 2 ms until atrioventricular (AV) block was first observed. Atrial arrhythmia (AA) was induced by 1 or 2 second bursts starting with a cycle length of 60 ms, decreasing successively with a 2-ms decrement, down to a cycle length of 10 ms. AA duration was the sum of time each mouse spent under and AA episode after completion of the two passes. Atrial arrhythmia (AA) inducibility was scored as the number of mice in which at least one episode lasting >1sec of arrhythmia was induced after pacing. Mice of both sexes were used for atrial arrhythmia induction experiments.

**Cellular electrophysiology**

Single cells were isolated from left atria of adult male mice by enzymatic dissociation. Therefore, excised hearts were perfused for 5 minutes in a Langendorff system with a modified Tyrode's solution containing (in mmol/L): NaCl 140, KCl 5.4, $CaCl_2$ 1.8, $MgCl_2$ 1.0, glucose 5.5, HEPES 5.0; pH 7.4 (set with NaOH). Subsequently, the hearts were perfused with Tyrode's solution containing a low $Ca^{2+}$-concentration (10 µmol/L) for 10 minutes, after which Liberase TM research grade (Roche Diagnostics, GmbH, Mannheim, Germany, 5401119001) and Elastase from porcine pancreas (Bio-Connect B.V., Huissen, Netherlands, W59168R) were added for 12 minutes at a concentration of 0.038 mg/mL and 0.01 mg/mL, respectively. All solutions were saturated with 100% $O_2$ and the temperature was maintained at 37 °C. To obtain single cells, the digested left atria was cut into small pieces which were triturated for 4 minutes through a pipette (tip diameter: 0.8 mm) in the low $Ca^{2+}$ Tyrode's solution, supplemented with 10 mg/ml Bovine Serum Albumin (Roche Diagnostics, essential fatty free, fraction V). Single cells were stored at room temperature for at least 45 min before they were used. Quiescent single rod-shaped cells with smooth surfaces were selected for electrophysiological measurements.

Action potentials (APs) were recorded with the amphotericin-B perforated patch-clamp technique using an Axopatch 200B amplifier (Molecular Devices Corporation, Sunnyvale, CA). Data acquisition and analysis were performed using custom software, and APs were low-pass filtered at 5 kHz and digitized at 40 kHz. Potentials were corrected for the estimated liquid junction potentials [10].

APs were recorded at 36±0.2°C using the modified Tyrode's solution. Pipettes (borosilicate glass (Harvard Apparatus, UK)) were filled with solution containing (in mmol/l): K-gluc 125, KCl 20, NaCl 5, amphotericin-B 0.44, HEPES 10, pH 7.2 (KOH). APs were elicited at 2 to 8 Hz by 3 ms, ~1.2× threshold current pulses through the patch pipette. We analyzed resting membrane potential (RMP), maximal AP amplitude (APA), maximum AP upstroke velocity, and AP duration at 20, 50 and 90% repolarization ($APD_{20}$, $APD_{50}$ and $APD_{90}$, respectively). Parameters from 10 consecutive APs were averaged.

**Intracellular $Ca^{2+}$ measurements**

Intracellular calcium concentrations: $[Ca^{2+}]_i$ were measured at 25°C in HEPES solution ((mmol/l): [Na+] 156, [K+] 4.7, [Ca2+] 1.3, [Mg2+] 2.0, [Cl−] 150.6, [HCO3] 4.3, [HPO 2−]

1.4, [Hepes] 17, [Glucose] 11 and 1% fatty acid free albumin, pH 7.3) using the fluorescent probe Indo-1 as described previously [11]. In brief, isolated myocytes were exposed to 5 µmol/l of the acetoxymethyl esters of indo-1 during 30 min at 37°C. Myocytes were attached to a poly-D-lysine (0.1 g/l) treated cover slip placed on a temperature-controlled microscope stage of an inverted fluorescence microscope (Nikon Diaphot) with quartz optics. A temperature-controlled perfusion chamber (height 0.4 mm, diameter 10 mm, volume 30 µL, temperature 37°C), with two needles at opposite sides for perfusion purposes, was tightly positioned over the cover slip. The contents of the chamber could be replaced within 100 ms. Bipolar square pulses for field stimulation (40 V/cm) were applied through two thin parallel platinum electrodes at a distance of 8 mm. One quiescent single myocyte was selected (myocytes with more than one spontaneous oscillation per 10 s were excluded) and the measuring area was adjusted to the cell surface with a rectangular diaphragm. The wavelength of excitation of Indo-1 was 340 nm, applied with a stabilized xenon-arc lamp (100 W). Fluorescence was measured in dual emission mode at 410 and 516 nm. Emitted light passed a barrier filter of 400 nm, a dichroic mirror (450 nm) and respective narrow band interference filters in front of two photomultipliers (Hamamatsu R-2949). Signals were digitized at 1 kHz and corrected for background signals recorded from Indo-1 free myocytes. Ten subsequent Ca2+ transients were averaged from which apparent $[Ca^{2+}]_i$ was calculated according to the ratio equation [12].

**Isolation of CM nuclei**

Nuclei isolation was performed as follows: snap frozen adult left and right atria from adult male and female mice were trimmed and homogenized in lysis buffer containing RNase inhibitor using an Ultra-Turrax homogenizer. Samples were further homogenized with a loose pestle douncer (10 strokes). After a 10-min incubation in the lysis buffer, an additional 10 strokes were performed with a tight pestle. The lysis procedure was monitored by light microscopy to ensure complete tissue and cell lysis and efficient nuclear extraction. The crude lysate was successively passed through 100  and 30µm mesh filters. The final lysate was spun at 1000 × g for 5 min and the resulting pellet was resuspended in 500 µl staining buffer (5% BSA in PBS) supplemented with RNAse inhibitor. Isolated nuclei were incubated with rabbit polyclonal antibodies specific for pericentriolar material 1 (PCM1) (Sigma-Aldrich; HPA023370) at a dilution of 1:400 for 1 h rotating at 4 °C. Next, Alexa Fluor 647-conjugated donkey-anti-rabbit 647 antibodies (ThermoFisher Scientific A-31573; 1:500 dilution), and DAPI (1:1000 dilution) were added and the incubation was continued for an additional hour. Samples were spun at 1000 × g for 10 min and washed with 500 µl staining buffer before resuspension in 500 µl staining buffer supplemented with RNAse inhibitor. Intact CM nuclei were sorted on a BD Influx FACS on the basis of DAPI and Alexa Fluor 647 positivity into cold BL+TG buffer from the ReliaPrep RNA Tissue Miniprep System (Promega, Z6112) for RNA isolation, and into resuspension buffer for ATACseq [13]. RNA was isolated following a gDNA depletion step according to the manufacturer's instructions. RNA yield and purity was assessed using an Agilent 2100 Bioanalyzer in combination with the RNA Pico chips.

### Library preparation and sequencing

For RNA isolated from left atria, 500 ng was used for library generation with the KAPA mRNA HyperPrep kit (Roche) and sequenced on the HiSeq4000 system (Illumina) with 50 bp single-end reads. RNAseq sample sizes are as follows: Whole adult left atria: 4 WT, 3 *RE(int)*$^{-/-}$, 3 *RE(down)*$^{-/-}$, 4 *Prrx1(enh)*$^{-/-}$, and 3 *Double homozygous*.

### Differential expression analysis

Reads were mapped to the mm10 build of the mouse transcriptome using STAR [14]. Differential expression analysis was performed using the DESeq2 package based on a negative binomial distribution model [15]. P-values were corrected for multiple testing using the false discovery rate (FDR) method of Benjamini-Hochberg. We have used 0.05 as FDR control level. Unsupervised hierarchical clustering was performed on differentially expressed genes using the R package pheatmap version 1.0.8. (http://cran.rproject.org/web/packages/pheatmap/index.html). PANTHER [16] was used for gene ontology (GO) biological process analysis. Benjamini–Hochberg correction was performed for multiple testing-controlled P values. Statistically significant enriched terms were functionally grouped and visualized.

### ATACseq on CM nuclei

ATACseq on FACS-sorted PCM1+ nuclei was performed and analyzed as described in [13]. Approximately 50k nuclei were used as input. The library was sequenced (paired-end 125 bp) and data was collected on a HiSeq4000.

### Peak-calling and motif analysis of ATACseq

Reads from ATAC-seq data were mapped to mm10 build of the mouse genome using BWA [17], the default settings were used. The BEDTools suite was used to distribute the genome wide signal into bins of 500 bp [18]. Bins with less than 89 cumulative tags across all 10 samples were discarded as noise. Differential accessibility was assessed using the DESeq2 package based on a model using the negative binomial distribution [15]. P-values were corrected for multiple testing using the false discovery rate (FDR) method of Benjamini-Hochberg. We have used 0.05 as FDR control level. Continuous bins with differential signal were subsequently merged together using the BEDTools suite.

200bp summits were determined of ATAC-seq performed in atrial cells [19]. In total, HOMER [20] was performed on sequences of 2,000 neutral called peaks (randomly sampled out of 85,570 peaks), 2,000 peaks up (randomly sampled out of 3,551 peaks), and 1,776 peaks down in *RE(int)*$^{-/-}$, all with random genome as background in HOMER.

Unsupervised hierarchical clustering was performed on differentially detected peaks and genes in RNA-seq and ATAC-seq using the R package pheatmap, version 1.0.8. (http://cran.r-project.org/web/packages/pheatmap/index.html).

### Usage of EMERGE

The genome-wide heart enhancer prediction track generated by EMERGE [21] was used as a proxy for the presence of putative heart enhancers. The particular prediction used aimed at identifying robustly active heart enhancers. This was accomplished by further annotating the available true positives [22] to train the algorithm with on the basis of the consistency in heart activity patterns shown. Full details are outlined in [4].

### Supplemental Figures

Supplemental figure 1

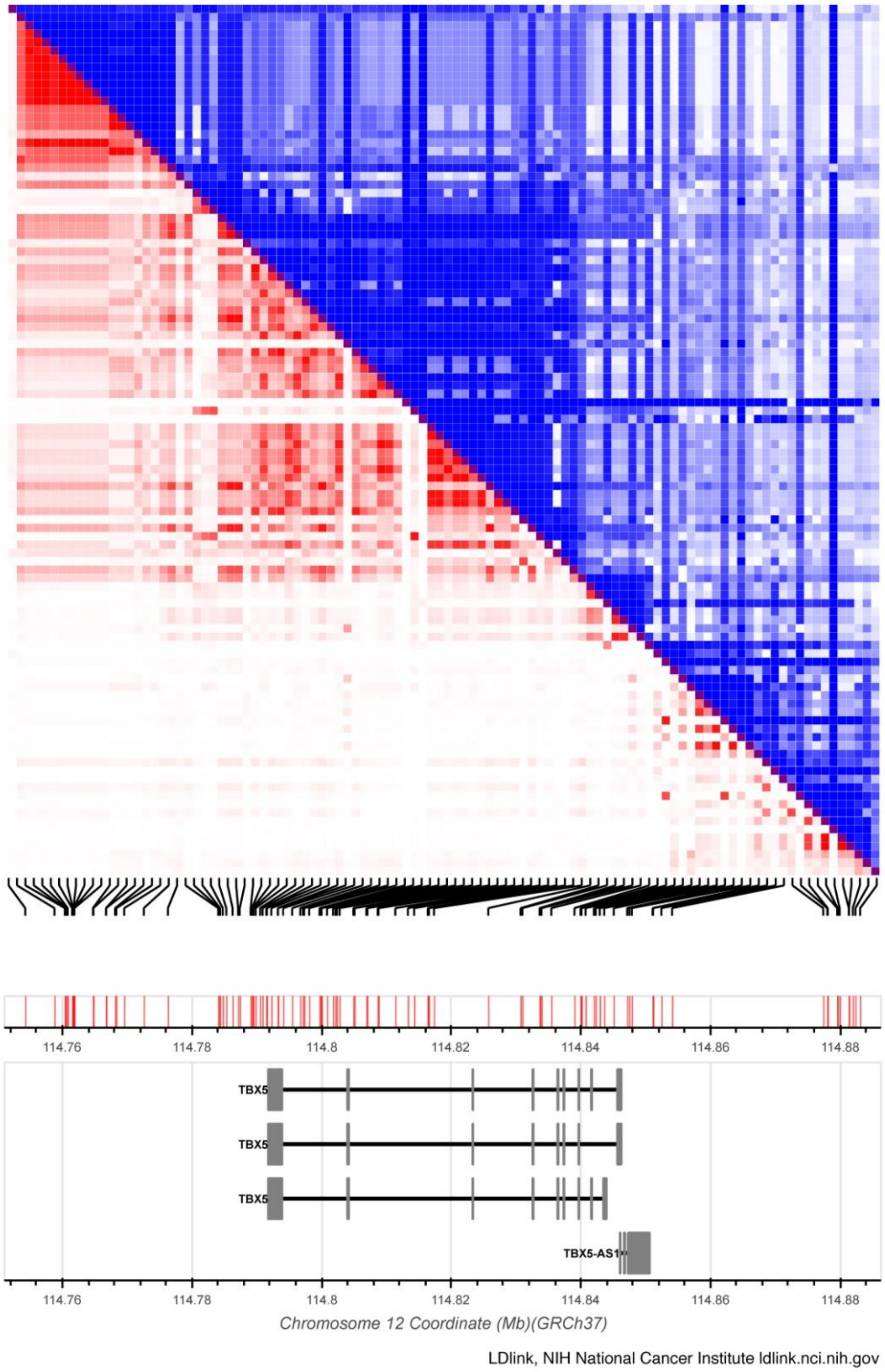


**Supplemental Figure 1. AF-associated variants near *TBX5* are grouped into distinct haplotypes.** LD matrix plot of SNPs located at the *TBX5* locus associated with AF. The SNPs found in the last intron and downstream of the gene are clustered into two distinct haplotypes based on minimum linkage disequilibrium r2 of 0.1. The plot was generated using the LD matrix tool al ldlink.nci.nih.gov.

Supplemental figure 2

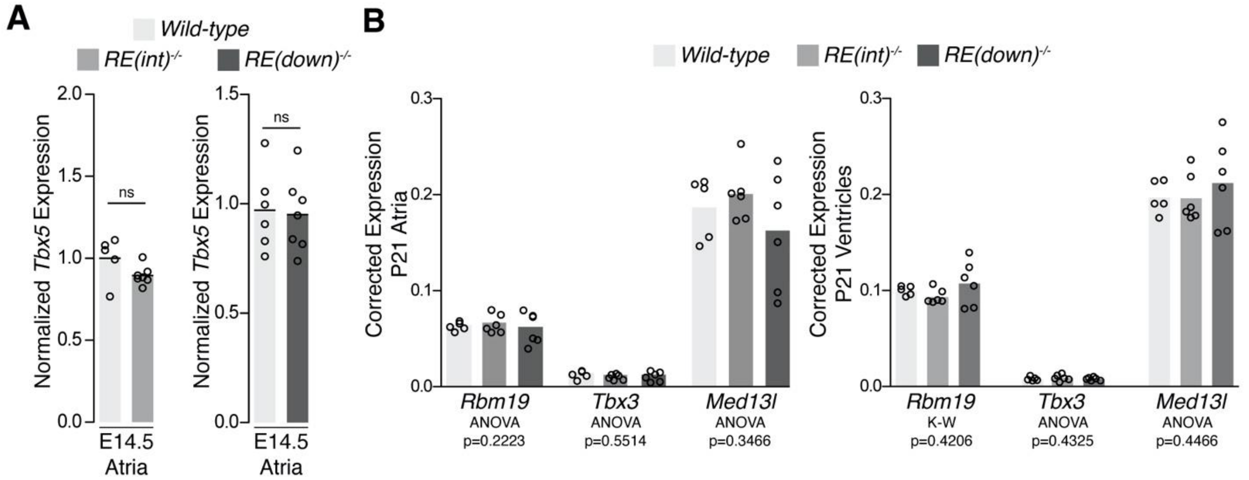


**Supplemental Figure 2. No difference in transcript levels of genes adjacent to *Tbx5* in deletion mutants.** A) Fetal (E14.5) expression levels of *Tbx5* were normalized to controls within each litter. B) Reference gene-corrected expression levels of *Rbm19*, *Tbx3*, and *Med13l* in juvenile whole atria (left) and ventricles (right) in *WT* (n=5), *RE(int)*$^{-/-}$ (n=6), and *RE(down)*$^{-/-}$ (n=6). Statistical significance within each gene and tissue type was determined with Kruskal-Wallis tests.

Supplemental figure 3

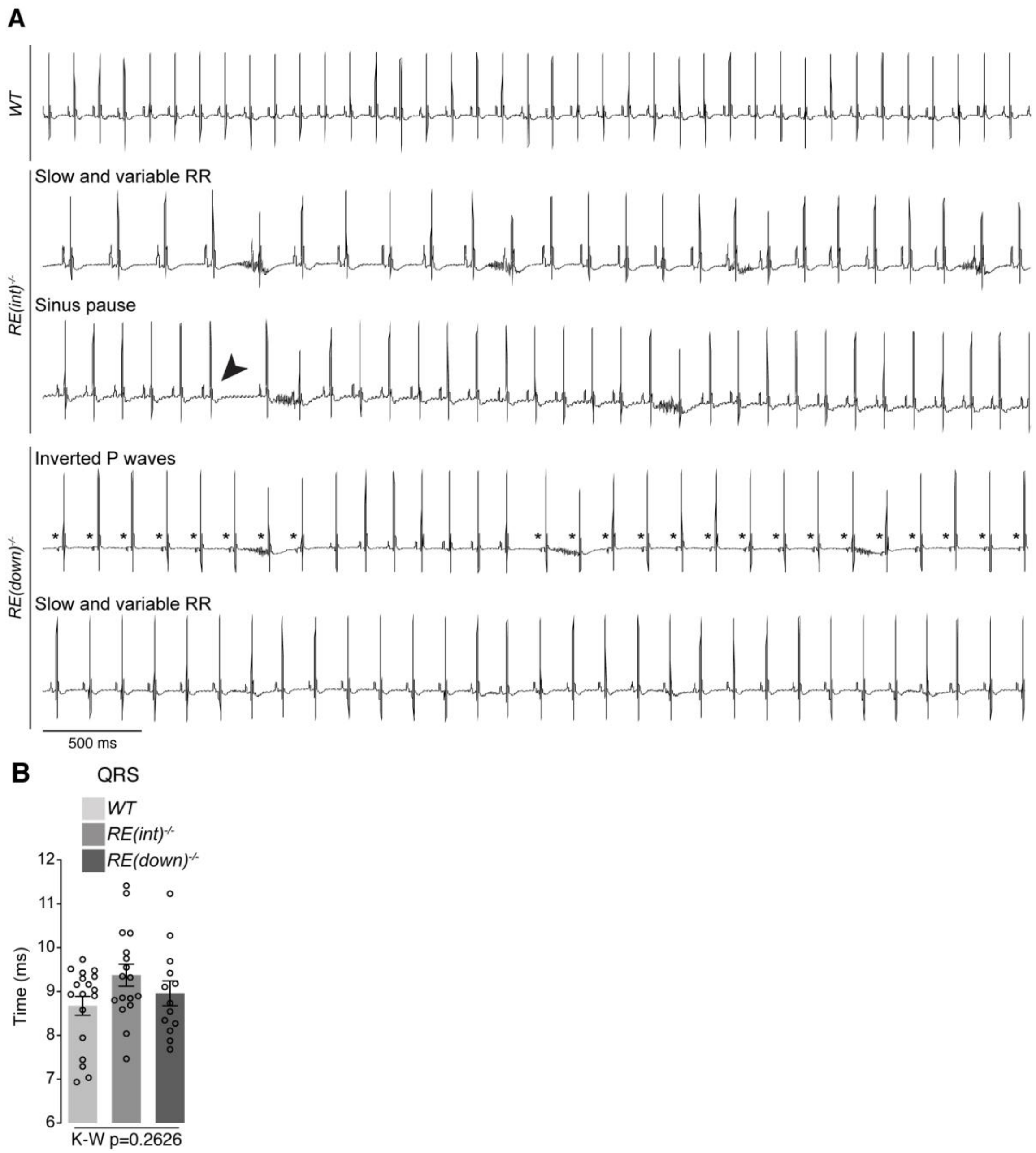


**Supplemental Figure 3. Additional *in vivo* electrophysiology parameters.** (A) Lead 2 traces of *WT* , *RE(int)*$^{-/-}$, and *RE(down)*$^{-/-}$ mice showing slow and variable RR, a sinus pause (arrowhead), or abnormal P waves (asterisk). (B) Graph shows no significant change in QRS duration in either mutant. Statistical significance was determined in B with Kruskal-Wallis test.

Supplemental figure 4

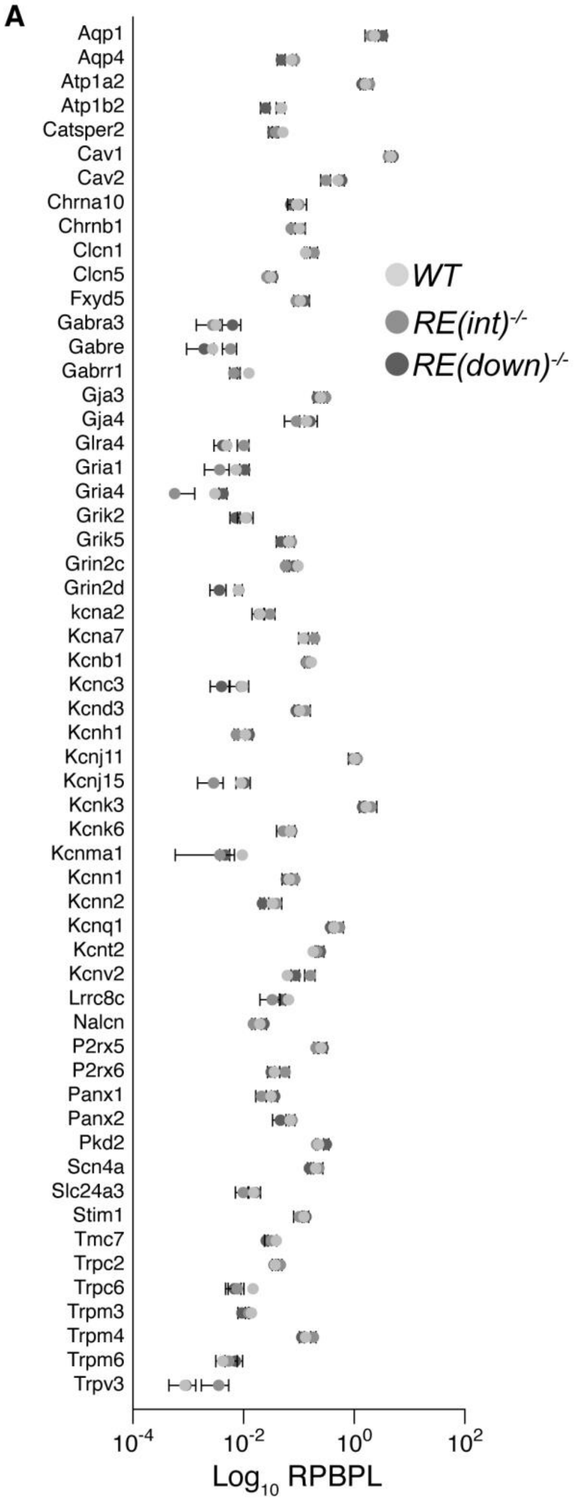


**Supplemental figure 4. Ion channel expression in left atria of deletion mutants.** (A) Graph depicts differential ion channel expression (fold change) in control and mutant left atria. RPBPL; Reads per (transcript) base pair length.

Supplemental figure 5

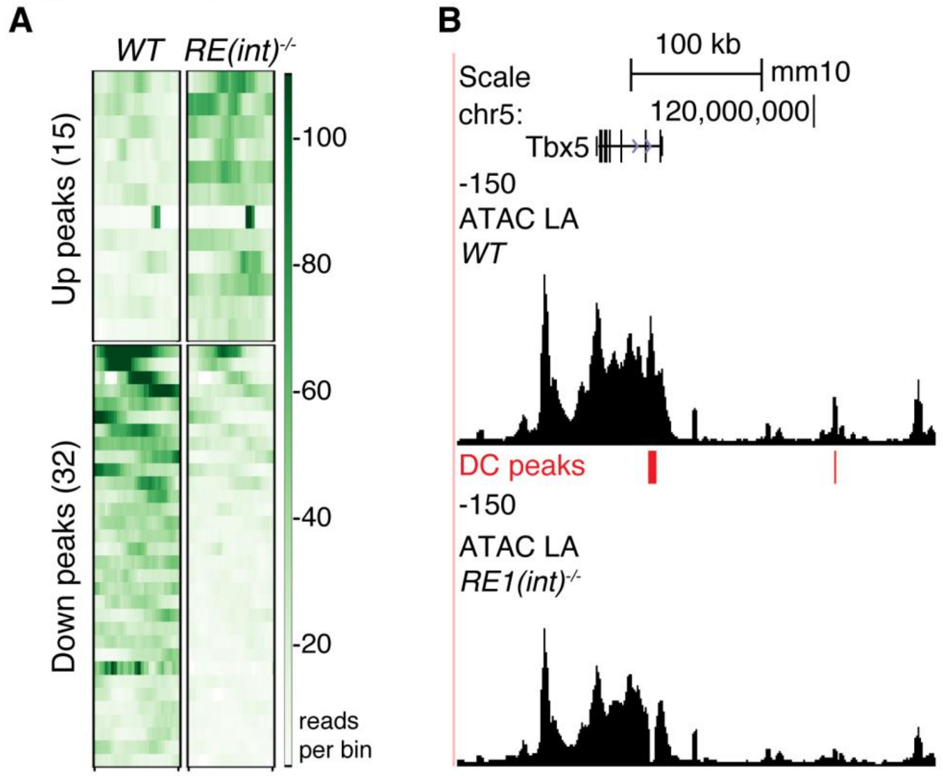


**Supplemental figure 5. Modest *Tbx5* upregulation minimally alters genome-wide chromatin accessibility.** (A) Heatmaps depicting vertically sorted up or downregulated accessible regions spanning 1500 bp bins (Padj<0.05) in *RE(int)*$^{-/-}$ (n=4) vs *WT* (n=4) cardiomyocytes. (B) UCSC browser view of the *Tbx5* locus showing differential accessibility profiles in mutants.

Supplemental figure 6

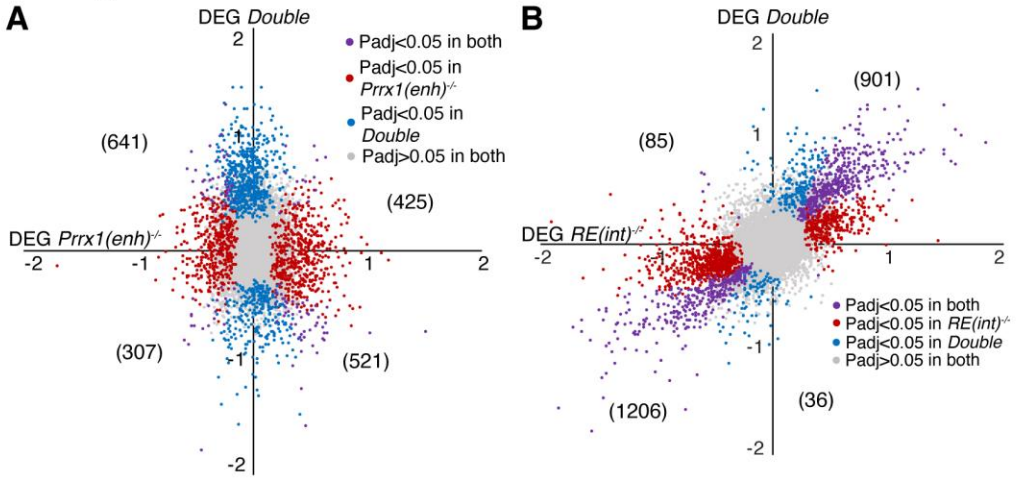


C

| GO biological process | *Prrx1(enh)*$^{-/-}$ UP/DOWN | Fold enrich | P-value | FDR | *RE1(int)*$^{-/-}$ UP/DOWN | Fold enrich | P-value | FDR | *RE1(int)*$^{-/-}$; *Prrx1(enh)*$^{-/-}$ UP/DOWN | Fold enrich | P-value | FDR |
|---|---|---|---|---|---|---|---|---|---|---|---|---|
| Regulation of biological quality (GO:0065008) | DOWN | 1.64 | 1.7E-08 | 3.04E-06 | UP | 1.55 | 1.52E-10 | 7.98E-07 | UP | 1.56 | 3.39E-08 | 1.49E-05 |
| Localization (GO:0051179) | DOWN | 2.11 | 1.47E-14 | 1.55E-11 | UP | 1.45 | 1.66E-09 | 6.54E-06 | UP | 1.44 | 2.87E-07 | 7.96E-05 |
| Developmental process (GO:0032502) | UP | 2.33 | 6.05E-20 | 1.59E-16 | UP | 1.39 | 6.75E-09 | 2.13E-05 | UP | 1.61 | 1.82E-14 | 2.88E-10 |
| Cellular component organization (GO:0016043) | DOWN | 1.85 | 4.25E-08 | 6.91E-06 | UP | 1.41 | 1.36E-08 | 3.06E-05 | UP | 1.54 | 1.28E-10 | 2.24E-07 |
| Regulation of ion transport (GO:0043269) | DOWN | 2.68 | 8.67E-08 | 1.32E-05 | UP | 2.22 | 1.69E-07 | 1.78E-04 | UP | 2.36 | 7.58E-07 | 1.79E-04 |
| Regulation of cell communication (GO:0010646) | DOWN | 2.18 | 2.84E-18 | 1.12E-14 | UP | 1.48 | 3.30E-07 | 2.90E-04 | UP | 1.57 | 3.36E-07 | 8.86E-05 |
| Positive regulation of cell projection organization (GO:0031346) | DOWN | 2.07 | 3.31E-04 | 1.58E-02 | UP | 2.44 | 1.46E-06 | 1.00E-03 | UP | 2.08 | 3.93E-07 | 1.00E-04 |
| Cardiac conduction (GO:0061337) | DOWN | 7.1 | 1.09E-04 | 5.97E-03 | UP | 6.57 | 3.60E-06 | 2.03E-03 | UP | 7.36 | 1.01E-05 | 1.50E-03 |
| Cell differentiation (GO:0030154) | UP | 2.25 | 2.84E-11 | 8.15E-09 | UP | 1.39 | 7.86E-06 | 2.76E-03 | UP | 1.65 | 4.77E-10 | 5.80E-07 |
| Metal ion transport (GO:0030001) | DOWN | 3.43 | 4.53E-08 | 7.30E-06 | UP | 2.18 | 1.02E-05 | 3.42E-03 | UP | 2.25 | 6.86E-04 | 4.90E-02 |
| Regulation of cell motility (GO:2000145) | UP | 2.96 | 2.10E-14 | 1.23E-11 | DOWN | 3.35 | 2.30E-38 | 9.10E-35 | DOWN | 2.28 | 2.53E-05 | 5.64E-03 |
| Extracellular matrix organization (GO:0030198) | UP | 7.57 | 1.96E-24 | 3.09E-20 | DOWN | 6.12 | 3.45E-34 | 4.19E-31 | DOWN | 7.87 | 3.22E-17 | 5.09E-13 |
| Circulatory system development (GO:0072359) | UP | 3.3 | 2.80E-17 | 3.69E-14 | DOWN | 3.2 | 3.27E-32 | 2.46E-29 | DOWN | 3.14 | 2.30E-10 | 9.09E-07 |
| Vasculature development (GO:0001944) | UP | 4.11 | 4.55E-17 | 4.79E-14 | DOWN | 3.97 | 1.63E-31 | 1.03E-28 | DOWN | 3.23 | 3.18E-07 | 1.74E-04 |
| Regulation of response to stimulus (GO:0048583) | UP | 1.91 | 1.84E-17 | 2.64E-14 | DOWN | 1.79 | 1.33E-28 | 5.83E-26 | DOWN | 1.61 | 2.18E-06 | 7.17E-04 |
| Actin cytoskeleton organization (GO:0030036) | DOWN | 3.17 | 4.57E-08 | 7.29E-06 | DOWN | 3.91 | 4.88E-28 | 1.88E-25 | DOWN | 3.26 | 7.89E-07 | 3.37E-04 |
| Cell adhesion (GO:0007155) | UP | 2.92 | 3.90E-12 | 1.50E-09 | DOWN | 3.1 | 1.83E-27 | 6.56E-25 | DOWN | 2.99 | 1.05E-08 | 1.85E-05 |
| Biological adhesion (GO:0022610) | UP | 2.94 | 1.97E-12 | 8.17E-10 | DOWN | 3.06 | 4.92E-27 | 1.62E-24 | DOWN | 2.95 | 1.41E-08 | 2.02E-05 |
| Angiogenesis (GO:0001525) | UP | 4.5 | 4.90E-12 | 1.84E-09 | DOWN | 4.87 | 4.94E-27 | 1.59E-24 | DOWN | 4.17 | 3.74E-07 | 1.91E-04 |
| Animal organ development (GO:0048513) | UP | 2.03 | 8.43E-17 | 7.39E-14 | DOWN | 1.84 | 1.25E-23 | 3.30E-21 | DOWN | 1.78 | 3.37E-07 | 1.77E-04 |

**Supplemental figure 6. Comparison of differentially expressed genes in *Prrx1(enh)*$^{-/-}$, *RE(int)*$^{-/-}$ , and *Double homozygous* adult left atria.** (A-B) X-Y plot of all transcripts in *Prrx1(enh)*$^{-/-}$ (A) or *RE(int)*$^{-/-}$ (B) **(**x axis) and *Double homozygous* (y axis), with deregulated genes (Padj<0.05) common to both genotypes in purple, *Prrx1(enh)*$^{-/-}$, or *RE(int)*$^{-/-}$ deregulated genes in red and *Double homozygous* deregulated genes in blue. (C) Gene ontology (GO) analysis of upregulated and downregulated genes in *RE(int)*$^{-/-}$ , *Prrx1(enh)*$^{-/-}$ , and *Double homozygous* samples.
